# Novel discovery of a lymphatic bridge connecting Schlemm’s canal to limbal and conjunctival lymphatic pathway

**DOI:** 10.1101/2023.04.26.538479

**Authors:** Yujia Yang, Meng Shi, Guangyu Li, Lejun Shen, Lu Chen

**Author notes:** **Corresponding author:** Lu Chen, MD and PhD, 689 Minor Hall, University of California, Berkeley, CA, 94720, USA;.

## Abstract

**Purpose:** Schlemm’s canal (SC) is a critical structure regulating aqueous humor (AH) drainage and intraocular pressure (IOP). It is known that in the conventional outflow pathway, AH flows from SC to episcleral veins. We recently reported a high-resolution three-dimensional (3D) imaging technology for intact eyeballs, SC and ocular surface. Using this advanced technology, we herein report the discovery of a new structure, termed lymphatic bridge, that directly connects SC to the limbal and conjunctival lymphatic pathway. Further investigation on this novel outflow pathway may provide new mechanisms and therapeutic approaches for glaucoma.

**Methods:** As reported previously, intact eyeballs were harvested from Prox-1-GFP (green fluorescent protein) mice and processed by a tissue clearing technique with CLARITY. Samples were immunolabeled with specific antibodies for CD31 (pan-endothelial marker) and LYVE-1 (lymphatic vessel endothelial hyaluronan receptor-1) and imaged by light-sheet fluorescent microscopy. The limbal areas were examined to locate connecting channels between SC and limbal and conjunctival lymphatic vessels. Moreover, in vivo anterior chamber dye injection was performed with Texas Red dextran for functional analysis on AH outflow.

**Results:** A novel lymphatic bridge structure that expressed both Prox-1 and LYVE-1 was discovered between the SC and limbal lymphatic vessels connected with conjunctival lymphatic pathway. Results from the anterior chamber dye injection showed AH drainage into the conjunctival lymphatic outflow pathway.

**Conclusions:** This study provides the first evidence on the direct connection between SC and limbal and conjunctival lymphatic pathway. This new pathway is distinctive from the conventional episcleral vein pathway and merits further investigation.

## 1. Introduction

Glaucoma is a major disease of irreversible blindness worldwide [1, 2]. It is a neurodegenerative disease with high intraocular pressure (IOP) as the primary risk factor. Schlemm’s canal (SC) is a critical structure regulating aqueous humor (AH) drainage and IOP. It is an important component of the conventional outflow pathway, which accounts for 70-90% of the total outflow in humans. The endothelial cell lining of SC is one of the primary sites of resistance to AH drainage and a major determinant of IOP. When SC resistance increases with age or under a pathological situation, IOP is elevated leading to optic nerve damage and vision loss. SC is therefore an important topic of investigation for glaucoma [3-10].

Previously, we reported the first evidence that SC expresses Prox-1 (prospero homeobox-1), the master control gene for lymphatic development [11]. Similar to lymphatic vessels, there are no red blood cells inside SC. SC does not express LYVE-1 (lymphatic vessel endothelial hyaluronan receptor-1), which is expressed after Prox-1 during development and is a marker for classical or typical lymphatic vessels. The lymphatic vessels on the ocular surface (including the limbus and conjunctiva) are classical lymphatic vessels that express both Prox-1 and LYVE-1. Although in glaucoma treatment, surgical or artificial channels are created to drain AH from the anterior chamber to subconjunctival space, there is no report on a natural or direct connection between the SC and ocular surface lymphatic vessels for AH drainage from the anterior chamber, which is the topic of our investigation.

Since SC a circumferential channel embedded at the iridocorneal angle of the anterior chamber, it is technically challenging to study this structure which is not accessible by conventional approaches. We have recently developed a high-resolution three-dimensional (3D) imaging technology for intact eyeballs, SC and ocular surface with tissue clearing technique with CLARITY (Clear Lipid-exchanged Acrylamide-hybridized Rigid Imaging/Immunostaining/*In situ* hybridization-compatible Tissue-hYdrogel) and light-sheet fluorescent microscopy [12]. This advanced technique provides the first chance for directly viewing of SC, its various surfaces as well as associated channels at the physiological status. The imaging processes can be performed with perspective, orthogonal, serial sectional, and rotational view, and the results can be presented in 3D images, videos, and surface rendered models. Using this powerful technology, we herein report the discovery of a novel structure, termed lymphatic bridge, that directly connects SC to the limbal and conjunctival lymphatic pathway. Moreover, like typical or classical lymphatic vessels on the ocular surface, this lymphatic bridge expresses both Prox-1 and LYVE-1. Our functional assay with anterior chamber dye injection has also confirmed AH drainage into this conjunctival lymphatic outflow pathway. This novel SC-lymphatic bridge-limbal and conjunctival lymphatic outflow pathway is distinctive from the traditional outflow pathway where AH drains from SC via the collector channels into episcleral veins, which are blood vessels. Further investigation on this novel phenomenon may offer new mechanisms and therapeutic approaches for glaucoma.

## 2. Materials and Methods

### 2.1. Animals

Adult (8-12 weeks) and postnatal (P) mice (P14) of Prox-1-GFP (green fluorescent protein) were used in this study, as previously reported [11, 13]. All animals were treated according to the ARVO Statement for the Use of Animals in Ophthalmic and Vision Research, and the protocols were approved by the Animal Care and Use Committee, University of California, Berkeley. A total of 34 mice were used in the study.

### 2.2. Tissue clearing

Intact eyeballs were fixed in 4% paraformaldehyde at 4°C for 24 hours and cleared as previously reported [12]. Briefly, fixed eyeballs were incubated in X-CLARITY™ hydrogel-initiator mixture solution at 4°C for 24 hours, and then polymerized at 37°C and -70kPa for 3 hours to form the tissue-hydrogel hybrids. The tissue-hydrogel hybrids were cleared by the X-

CLARITY electrophoretic system (Logos Biosystems, Annandale, VA, USA) at 37°C with 1.0 A current and 50 rpm pump speed for 20 hours. Cleared samples were washed thoroughly in phosphate-buffered saline (PBS) and stored at 4°C for further processing.

### 2.3. Immunolabeling

Cleared samples were labeled by CD31 (goat anti-mouse, R&D, Minneapolis, MN, USA) or lymphatic endothelial marker LYVE-1 (rabbit anti-mouse, Abcam, Cambridge, MA, USA) with the DeepLabel antibody labeling kit (Logos Biosystem, Annandale, VA, USA) following the manufacturer’s instruction. Samples were sequentially incubated in the primary antibodies at 37°C for 3 days, washed in PBS for 6 hours, and incubated in the secondary antibodies at 37°C for 3 days.

### 2.4. Light sheet microscopy

For cleared samples, the refractive index was matched by immersing in the X-CALRITY mounting solution before imaging with a light sheet fluorescent microscopy (Zeiss Lightsheet Z.1, Carl Zeiss AG, Germany). Laser lines of 488-nm and 561-nm, Illumination Optics Lightsheet Z.1 5×/0.1 were used for illumination, and Detection Optics Lightsheet Z.1 5×/0.16 were used for detection. Stack images were taken along the Z or X axis (Figure 1A, upper left panel). 3D images and videos were generated by Bitplane Imaris x64 software, version 9.3.0 (Bitplane AG, Zurich, Switzerland).

**Figure 1.**
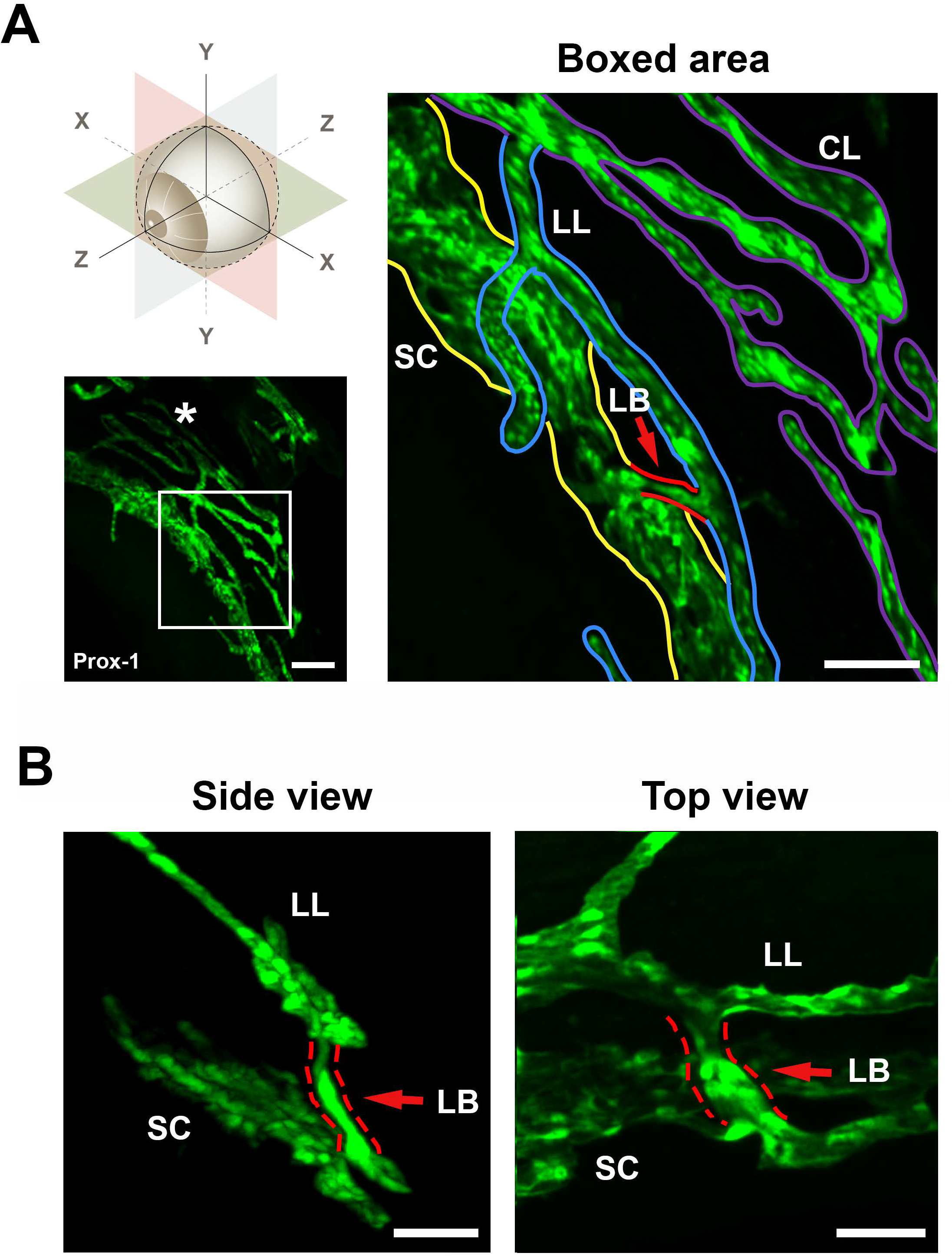
Representative images showing the lymphatic bridge between the Schlemm’s canal and limbal lymphatic vessels. **A**. The anterior segment of an adult Prox-1-GFP mouse was imaged along the Z-axis demonstrated in the upper left panel. Lower left panel, Prox-1^+^ vascular structures (green) were detected at the Schlemm’s canal and ocular surface. White asterisk: nictating membrane. Right panel, the enlarged boxed area showing the location of the lymphatic bridge (LB, red arrow and outlines) between the Schlemm’s canal (SC, yellow outlines) and limbal lymphatic vessels (LL, blue outlines), which were continuously connected with conjunctival lymphatics (CL, purple outlines). **B**. Side (left) and top (right) view from 3D imaging showing the lymphatic bridge (LB, red arrows and outlines) between Schlemm’s canal (SC) and limbal lymphatic vessels (LL). Scale bars: 100 μm (A), 50 μm (B).

### 2.5. Anterior chamber dye injection and intravital imaging

The experiment was performed using our advanced live imaging system as previously reported [11, 13, 14]. Mice were anesthetized with isoflurane and topical proparacaine hydrochloride ophthalmic solution (0.5%, Sandoz Inc, Princeton, NJ, USA). The cornea was penetrated with a 34G needle connected with a 10 μl Hamilton syringe (Hamilton, Reno, NV, USA). Fluorescently labeled dye (Texas Red™ Dextran, Invitrogen, USA) were injected into the anterior chamber.

Images were taken using a customized imaging system consisting of Zeiss Axio Zoom V.16 (Carl Zeiss AG, Gottingen, Germany) [11, 13, 14]. The captured Z-stack images were processed with Helicon Focus imaging software (Heliconsoft Ltd., http://www.heliconsoft.com) to obtain extended focus.

## 3. Results

### 3.1. A novel lymphatic bridge structure was identified between Schlemm’s canal and limbal lymphatic vessels

We first examined the anterior segments of adult mouse eyeballs with the ring of SC at the XY plane and tracked the Prox-1^+^ exiting routes from the SC. A novel Prox-1^+^ structure, termed lymphatic bridge, was discovered among the samples and this structure was observed to directly connect the SC with limbal lymphatics. As shown in Figure 1 and Supplementary Video 1 and 2, the entire route of the SC-lymphatic bridge-limbal lymphatics is Prox-1 positive, as well as the conjunctival lymphatics that are continuously connected with the limbal lymphatics.

### 3.2. The lymphatic bridge is distinctive from the collector channel

We further differentiated the lymphatic bridge from collector channels by immunostaining with CD31, the pan-endothelial marker. The collector channels belong to the conventional outflow pathway that drains into the episcleral veins, which are blood vessels and Prox-1 negative. As shown in Figure 2, two distinctive outflow pathways were identified. While the lymphatic bridge was connected to the Prox-1^positive^ CD31^positive^ limbal lymphatics, the collector channel was linked to the Prox-1^negative^ CD31^positive^ blood vessels belonging to the episcleral vein pathway.

**Figure 2.**
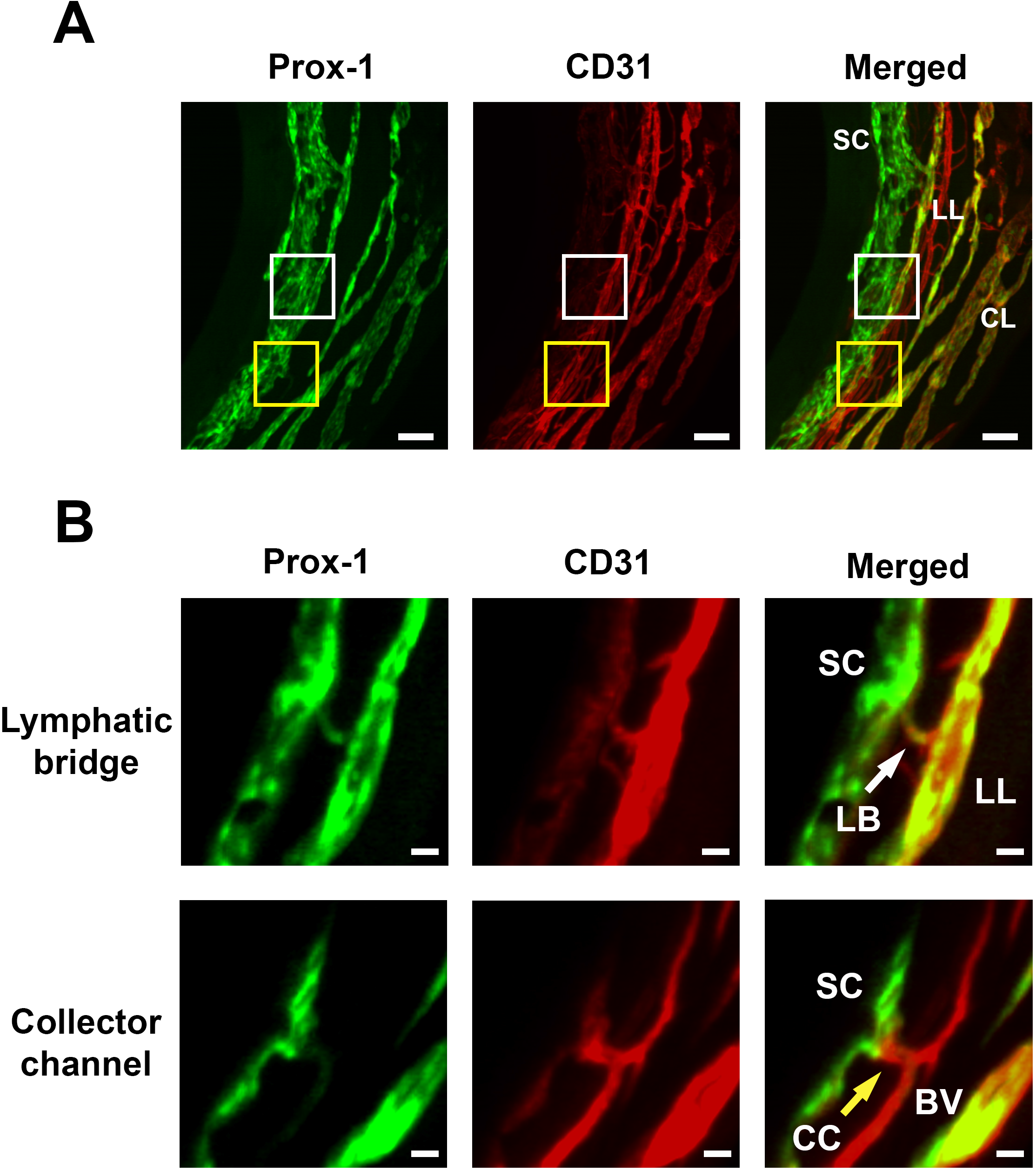
Representative images showing two distinctive outflow pathways from the Schlemm’s canal. Images were taken at the limbal area of an adult Prox-1-GFP mouse eyeball labeled with CD31. **A**. Top view of 3D imaging of the limbal area. The stacked image shows various limbal vascular structures detected including Schlemm’s canal (SC), limbal lymphatics (LL), and conjunctival lymphatics (CL). The white and yellow boxed area indicate the locations of the lymphatic bridge and collector channel, respectively. **B**. Extended sectional views from the boxed areas in panel A. Upper panels: the lymphatic pathway where the lymphatic bridge (LB, white arrow) connected the Schlemm’s canal (SC) with the Prox-1^positive^ CD31^positive^ limbal lymphatics (LL). Lower panels: the blood vascular pathway where the collector channel (CC, yellow arrow) linked the Schlemm’s canal (SC) to the Prox-1^negative^ CD31^positive^ blood vessel (BV) belonging to the episcleral vein pathway. Prox-1: green; CD31, red. Scale bars: 200 μm (A), 20 μm (B).

### 3.3. The lymphatic bridge was present during developmental stage

We next examined whether the lymphatic bridge is present during development. It is known that SC develops from flattened core of cells to discontinuous lumen by P14 [15]. As shown in Figure 3 and Supplementary Video 3, the Prox-1^positive^ CD31^positive^ lymphatic bridge structure was also detected among the P14 eyeballs in development.

**Figure 3.**
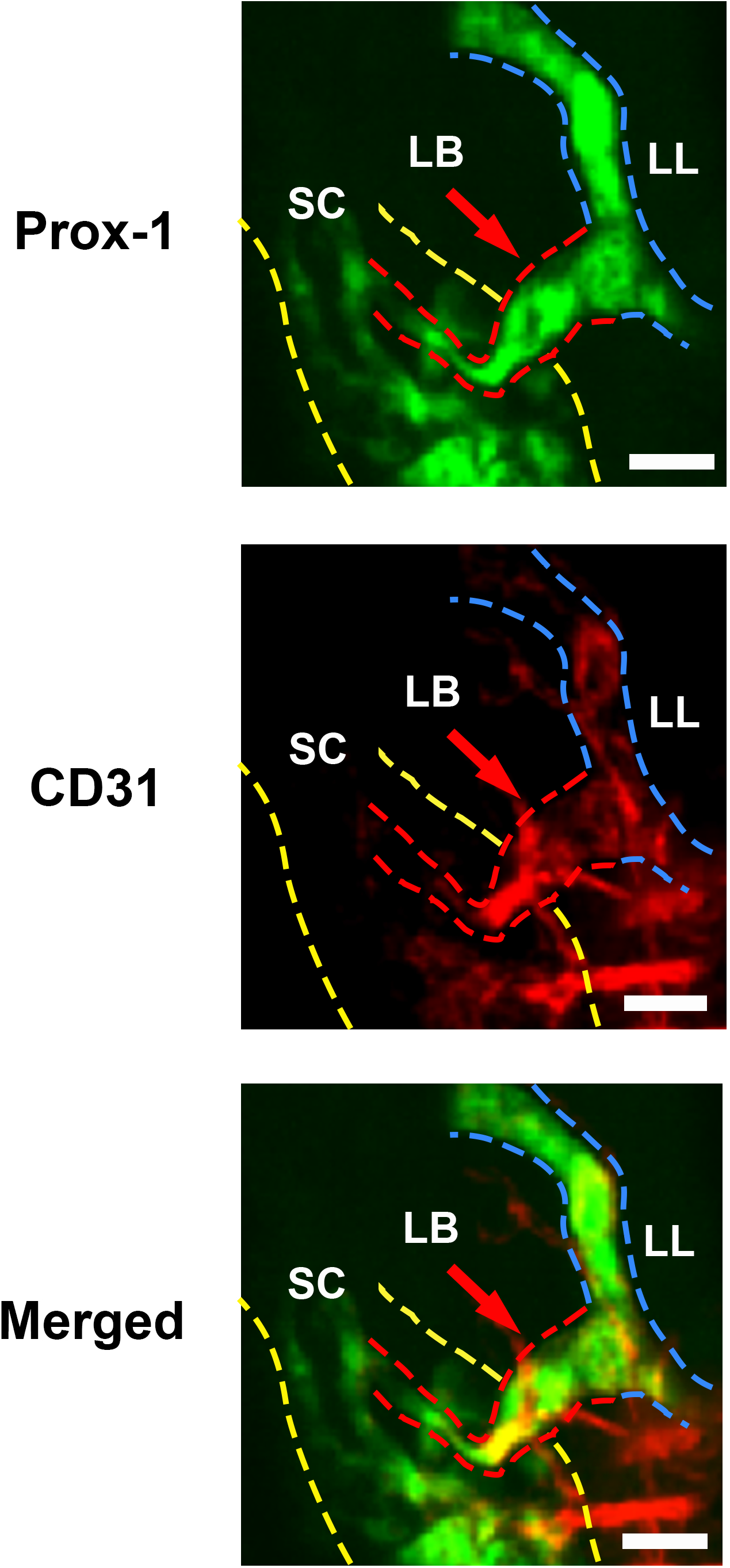
Representative images showing the presence of lymphatic bridge in development. Images were taken at the limbal area of a P14 Prox-1-GFP mouse eyeball labeled with CD31 and presented in the extended sectional view. The lymphatic bridge (LB, red arrows and red outlines) was detected between the Schlemm’s canal (SC, yellow outlines) and the limbal lymphatics (LL, blue outlines). Prox-1: green; CD31, red. Scale bars: 30 μm.

### 3.4. The lymphatic bridge expressed LYVE-1

Next, we examined the eyeballs with LYVE-1, the widely used lymphatic marker for classical or typical lymphatic vessels. As shown in Figure 4, it was found that the Prox-1^+^ lymphatic bridge co-expressed LYVE-1, confirming it is a classical or typical lymphatic structure.

**Figure 4.**
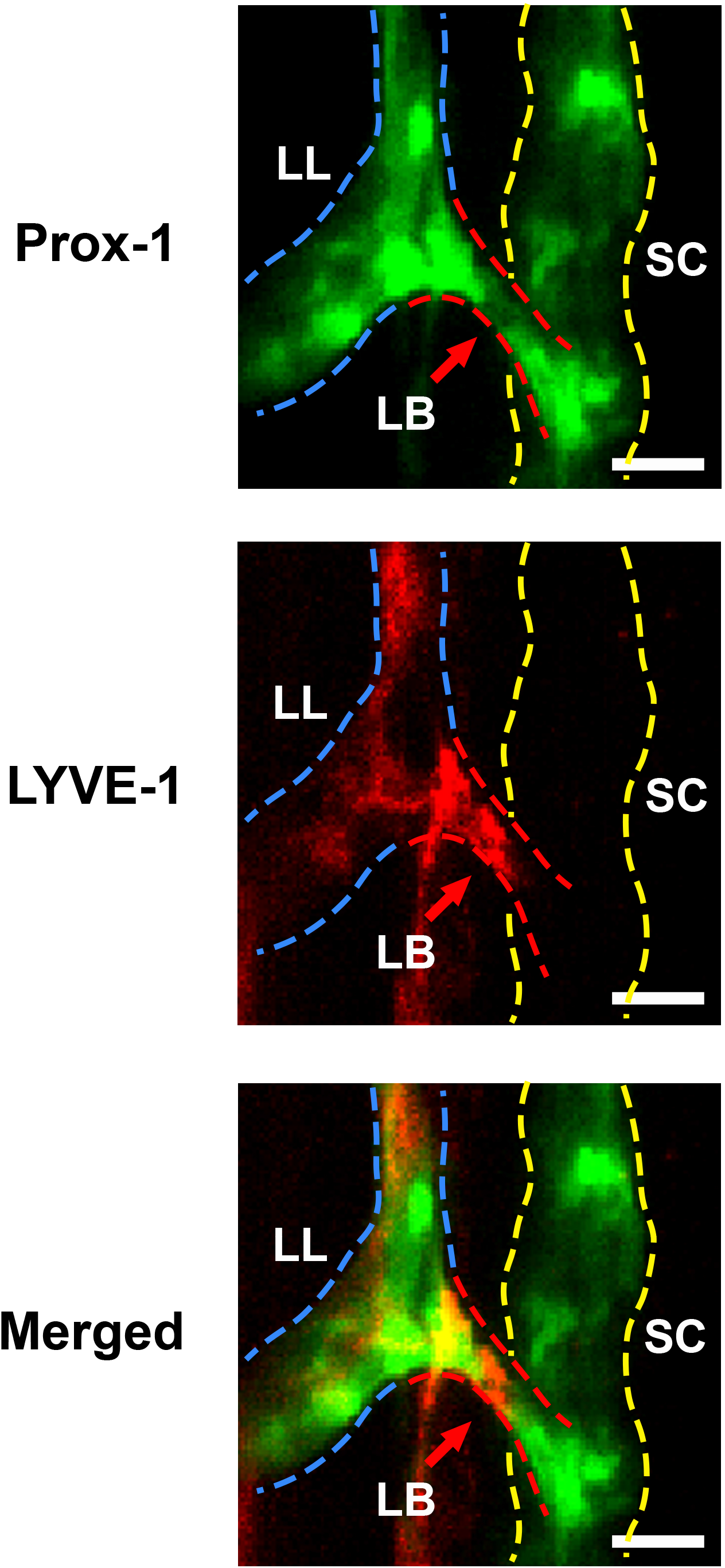
Representative images showing the lymphatic bridge expressed both Prox-1 and LYVE-1. Images were taken from an adult Prox-1-GFP mouse eyeball labeled by LYVE-1. The Prox-1^positive^ LYVE-1^positive^ lymphatic bridge (LB, red arrows and red outlines) was detected between the Prox-1^positive^ LYVE-1^negative^ Schlemm’s canal (SC, yellow outlines) and Prox-1^positive^ LYVE-1^positive^ limbal lymphatics (LL, blue outlines). Similar to limbal lymphatics, the lymphatic bridge expressed both Prox-1 and LYVE-1. Prox-1: green; LYVE-1, red. Scale bars: 20 μm.

### 3.5. Aqueous humor drained into the conjunctival lymphatic pathway from the anterior chamber

We further performed a functional study of anterior chamber dye injection and live imaging of AH outflow using our advanced live imaging system. As shown in Figure 5, the fluorescent signal was detected inside the Prox-1^+^ conjunctival lymphatic vessels after the injection of Texas Red™ Dextran into the anterior chamber. This study revealed a novel conjunctival lymphatic outflow pathway for AH drainage from the anterior chamber.

**Figure 5.**
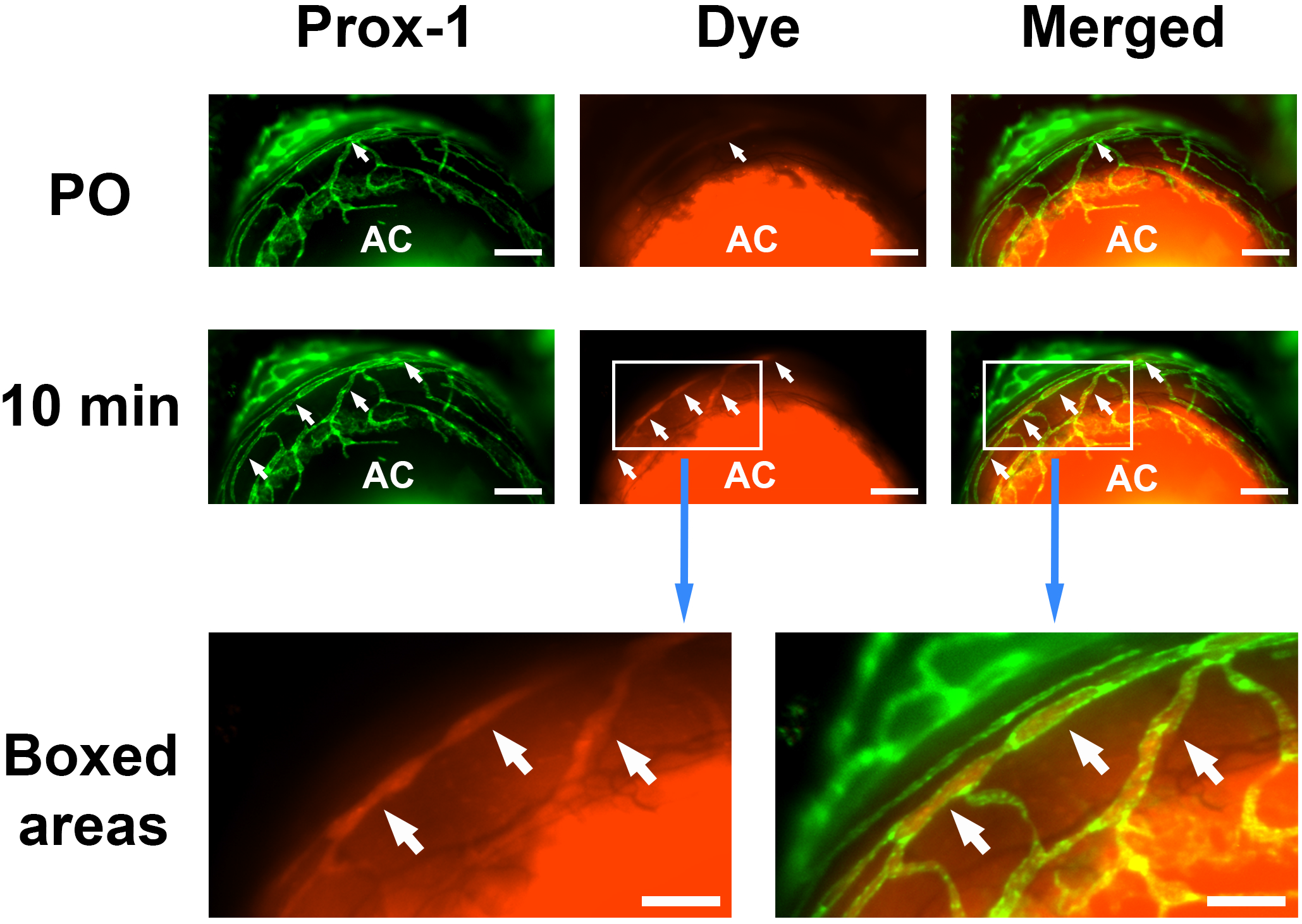
Representative images showing the drainage of fluorescent dye into the conjunctival lymphatic vessels after anterior chamber injection. Intravital images showing that the Texas Red fluorescent dye (red) injected into the anterior chamber (AC) was detected inside the Prox-1^+^ conjunctival lymphatics (green) post-operation (PO, upper panels) and 10 minutes after the injection (middle panels), as indicated by the white arrows. Bottom panels: enlarged images from the boxed areas. Scale bars: 200 μm (PO, 10 minutes), 100 μm (boxed areas).

## 4. Discussion

In this study, we report a novel lymphatic bridge structure that directly links SC with the limbal and conjunctival lymphatic pathway on the ocular surface. The study challenges the traditional view about the conventional outflow pathway in that AH drains from SC into the episcleral veins, which are blood vessels. It has been considered that there is no direct connection between the SC and lymphatic vessels. As illustrated in Figure 6, compared with the episcleral vein pathway which only expresses Prox-1 at the initial segment around SC, the entire conjunctival outflow pathway (SC-lymphatic bridge-limbal and conjunctival lymphatic vessels) are Prox-1 positive. Moreover, starting from the lymphatic bridge, all structures involved in this new conjunctival outflow pathway also expresses LYVE-1 indicating they are classical or typical lymphatic vessels. Since the conjunctiva is endowed with a rich network of lymphatics [16], this study reveals an important contributor for AH drainage from the SC. Given that a lymphatic feature is also reported in the unconventional pathway via uveoscleral outflow [17, 18], the lymphatic system taken together plays a more significant role in AH drainage than previously recognized.

**Figure 6.**
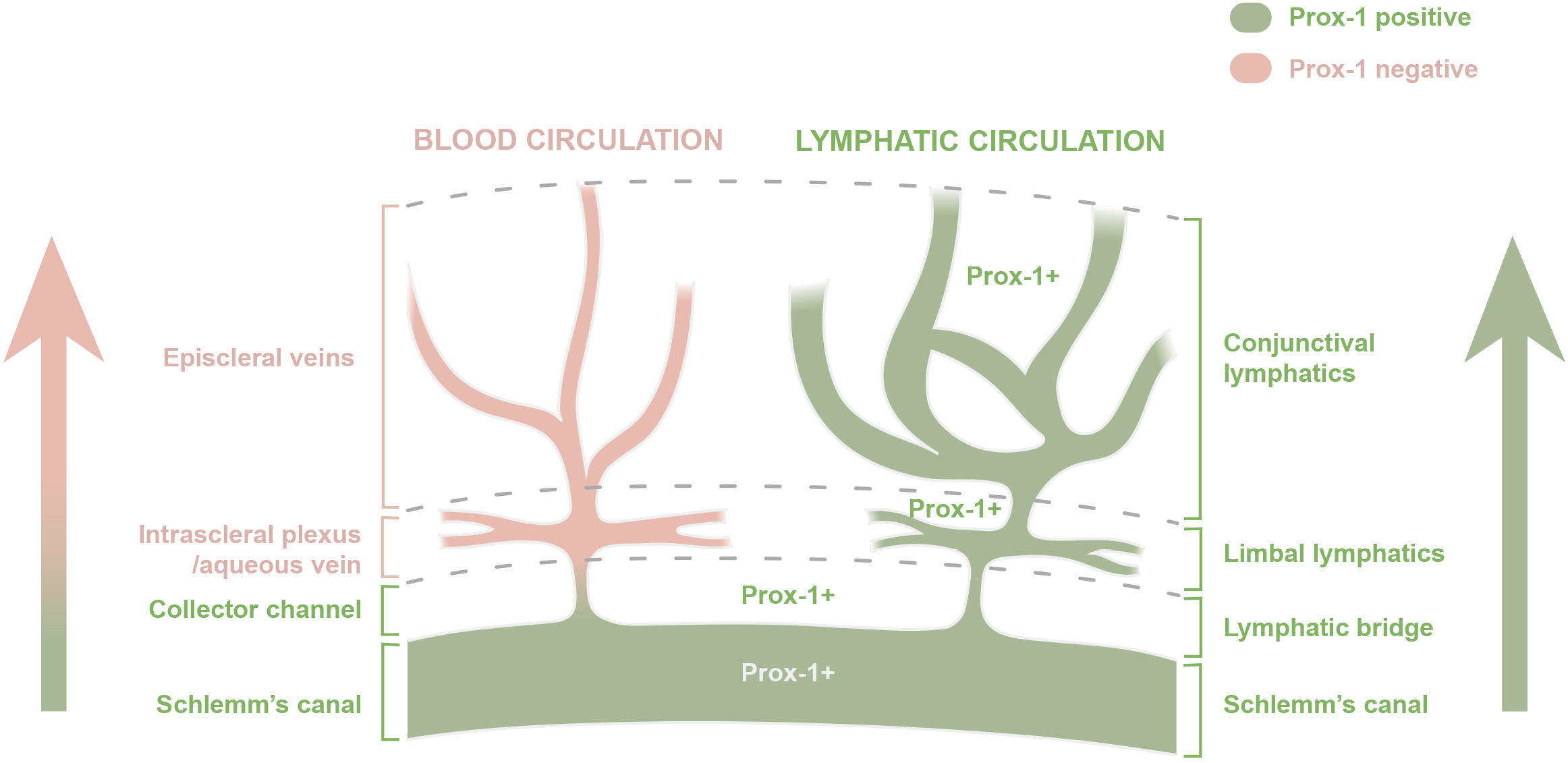
Schematic diagram showing the difference between the conjunctival lymphatic outflow pathway and the episcleral outflow pathway. Left, the episcleral vein pathway where aqueous humor flows from the Schlemm’s canal to collector channels, intrascleral plexus/aqueous vein, and episcleral veins, which are blood vessels. Right, the novel conjunctival lymphatic outflow pathway where aqueous humor drains from Schlemm’s canal to lymphatic bridge, and limbal and conjunctival lymphatic vessels. In the episcleral pathway, only the initial segment around the Schlemm’s canal expresses Prox-1, while the entire conjunctival outflow pathway is Prox-1 positive. Structures of the conjunctival outflow pathway are typical lymphatic vessels that express both Prox-1 and LYVE-1.

On the ocular surface, in contrast to the cornea which is by nature devoid of lymphatic vessels, the conjunctiva is rich in lymphatic supply [16]. Conjunctiva is a common site involved in glaucoma treatment, such as with the filtration surgeries where artificial channels are created to drain AH into the subconjunctival space from the anterior chamber. Our knowledge on conjunctival lymphatics, AH drainage, and glaucoma is rather limited at this stage. A recent study with post-mortem eyeballs shows that conjunctival lymphatics can drain from artificial blebs created by subconjunctival injection of fluorescent tracers [19]. However, in this study, the blebs are created locally at the conjunctiva and they are not connected to the anterior chamber.

Another recent study on post-trabeculectomy patients shows that eyes with conjunctival lymphatic connections to subconjunctival blebs have lower IOP compared to those without lymphatic connections [20]. However, surgical procedures with artificial channels are invasive and often fail with complications, such as scar formation. It is therefore plausible to facilitate AH drainage via the new natural conjunctival lymphatic pathway that directly drains from the anterior chamber. This merits further investigation.

In this study, we noticed that some of the collector channels express Prox-1, which is consistent with a previous report [21]. The lymphatic bridge can be distinguished from the collector channels by its connection with the Prox-1 positive limbal lymphatic vessels. In contrast, the collector channels are linked to the Prox-1 negative blood vessels that belong to the episcleral vein pathway. We didn’t observe lymphatic bridge in all samples. This may be due to individual variation or experimental limitation. It may also explain the differential propensity to glaucoma among populations, which warrants further investigation as well.

In conclusion, this study reports a novel lymphatic bridge structure that directly connects SC with the conjunctival lymphatic outflow pathway on the ocular surface. This pathway consisting of classical lymphatic vessels is distinctive from the traditional episcleral vein pathway draining into blood vessels. It is anticipated that further investigation on the lymphatic drainage system may provide novel mechanisms and therapeutic strategies to manage AH outflow and glaucoma in the future.

## Acknowledgements

This work is supported in part by research grants from the National Institutes of Health and the University of California at Berkeley (L.C.). The funding sources had no role in study design, the collection, analysis and interpretation of data, the writing of the report, or the decision to submit the article for publication. We thank Young K. Hong at University of Southern California and the Mutant Mouse Regional Resource Centers (MMRRC) for providing the founder Prox-1 transgenic mice. Lightsheet microscope imaging was conducted at the CRL Molecular Imaging Center, Helen Wills Neuroscience Institute at the University of California at Berkeley.

## Supplementary Videos

**Supplementary video 1**. Top view showing continuous vascular structures from the Schlemm’s canal (SC) to lymphatic bridge (LB, red arrow) and limbal lymphatic vessel (LL). Video was taken from continuous cross-sectional imaging of the limbal area of an adult Prox-1-GFP (green) mouse.

**Supplementary video 2**. Fluorescent 3D image of the same lymphatic bridge (LB, red arrow) shown in Supplementary video 1, rotated to show structural connections with the Schlemm’s canal (SC) and limbal lymphatics (LL). Green, Prox-1.

**Supplementary video 3**. Fluorescent 3D image of the lymphatic bridge (LB, red arrow) detected in a P14 Prox-1-GFP (green) mouse eyeball. SC: Schlemm’s canal, LL: limbal lymphatics.

